# Macroscale estimates of species abundance reveal evolutionary drivers of biodiversity

**DOI:** 10.1101/426379

**Authors:** Keiichi Fukaya, Buntarou Kusumoto, Takayuki Shiono, Junichi Fujinuma, Yasuhiro Kubota

## Abstract

Evolutionary processes underpin the biodiversity on the planet. Theories advocate that the form of the species abundance distribution (SAD), presented by the number of individuals for each species within an ecological community, is intimately linked to speciation modes such as point mutation and random fission. This prediction has rarely been, however, verified empirically; the fact that species abundance data can be obtained only from local communities critically limits our ability to infer the role of macroevolution in shaping ecological patterns. Here, we developed a novel statistical model to estimate macroscale SADs, the hidden macroecological property, by integrating spatially replicated multispecies detection-nondetection observations and the data on species geographic distributions. We determined abundance of 1,248 woody plant species at a 10 km grid square resolution over East Asian islands across subtropical to temperate biomes, which produced a metacommunity (i.e. species pool) SAD in four insular ecoregions along with its absolute size. The metacommunity SADs indicated lognormal-like distributions, which were well explained by the unified neutral theory of biodiversity and biogeography (UNTB) with protracted speciation, a mode of speciation intermediate between point mutation and random fission. Furthermore, the analyses yielded an estimate of speciation rate in each region that highlighted the importance of geographic characteristics in macroevolutionary processes and predicted the average species lifetime that was congruent with previous estimates. The estimation of macroscale SADs plays a remarkable role in revealing evolutionary diversification of regional species pools.

---

A better understanding of global patterns of species commonness and rarity has been a fundamental requirement in ecology and evolutionary biology since the time of Darwin (1859) (Hutchinson 1959, May 1988, Rosenzweig 1995). Nonetheless, we still lack a clear understanding of the patterns of species abundance, especially at large spatial scales, such as those representing regional species pools. The unified neutral theory of biodiversity and biogeography (UNTB; Hubbell 2001) provides a mechanistic explanation of the origin and maintenance of biodiversity; based on the premise that all individuals in a system are functionally equivalent and thus follow neutral processes of demography, dispersal, and speciation, the UNTB derives species abundance distributions (SADs), at both local-community and meta-community (i.e. species pool) scales, in addition to a range of other macroecological and macroevolutionary patterns such as the species-area relationship (Rosindell *et al*. 2011), *β* diversity (Chave & Leigh 2002), and various phylogeny characteristics (Davies *et al*. 2011).

The UNTB bridges evolutionary biology and community ecology by linking, theoretically, macroevolutionary processes to biodiversity patterns. In particular, it predicts that the statistical form of the SAD in the metacommunity is dependent on the mode of speciation (Hubbell 2001, Etienne *et al*. 2007, Haegeman & Etienne 2010, Rosindell *et al*. 2010, Etienne & Haegeman 2011, Haegeman & Etienne 2017). The point mutation speciation model, which formed the basis of the first UNTB proposed by Hubbell (2001), models speciation as a process in which each new species is represented initially by a single individual. The point mutation speciation model predicts a metacommunity SAD that follows the logseries distribution, a distribution that is characterized by a relatively high proportion of rare species (Hubbell 2001, Etienne & Alonso 2005). In contrast, the random fission speciation model assumes that speciation occurs in the metacommunity owing to the random division of a population of an existing species. The random fission speciation model predicts a fairly even metacommunity structure, which is related to the MacArthur’s (1957) broken-stick model (Haegeman & Etienne 2010, Etienne & Haegeman 2011). The point mutation speciation and random fission speciation represent the two extremes of a spectrum of speciation modes in UNTB. This spectrum of speciation modes has been argued to be unified with the concept of protracted speciation, which characterizes speciation as a gradual, drawn-out process (Rosindell *et al*. 2010, Haegeman & Etienne 2017). The UNTB with protracted speciation predicts a metacommunity SAD that follows a difference-logseries distribution. The difference-logseries distribution follows a logseries distribution at large abundances while behaving differently at small abundances; namely, it predicts fewer rare species than the logseries distribution (Rosindell *et al*. 2010, Haegeman & Etienne 2017).

Our ability to infer evolutionary processes that underpin observed biodiversity patterns is, however, fundamentally limited because species abundance data can be obtained only from local communities. Indeed, earlier studies have shown that differences in the mode of speciation are hardly discerned based on samples from local communities as they may not leave a signature on SADs realized in dispersal-limited localities (Hubbell 2001, Etienne *et al*. 2007, Rosindell *et al*. 2010, Etienne & Haegeman 2011). The limitation in data acquisition also prohibits us from identifying the rate of speciation (*ν*) from SADs because local community SADs are determined by the fundamental biodiversity number (*θ*), which is a compound parameter depending both on *ν* and the metacommunity size (*J_M_*) (Etienne & Alonso 2005, Etienne & Haegeman 2011; but see Etienne *et al*. 2007). Consequently, fundamental macroevolutionary properties of a metacommunity, such as *ν* and the average lifespan of the species (*L*; Ricklefs 2003), have remained largely unknown.

A solution to these problems is to obtain data on species abundance over a huge spatial extent that directly informs about the size and biodiversity of the metacommunity; such data is, however, unrealistic. In this view, we developed a novel hierarchical model (Royle & Dorazio 2008, Kéry & Schaub 2012, Kéry & Royle 2016) that estimates SADs over a large geographic extent, which we named “macroscale SADs”. The model integrates spatially replicated multispecies detection-nondetection observations and information on the geographical distribution of species. We applied the model to a large dataset of woody plant communities in midlatitude forests on East Asian islands, including the Japanese archipelago. The dataset comprised more than 40 thousand vegetation survey records and various data sources for geographical ranges of species. The model enabled us to estimate macroscale abundance for 1,248 species at a 10 km grid square resolution.

Although defining a metacommunity is difficult in practice, discerned biogeographic divisions will proximate its theoretical definition as they can be regarded an evolutionary unit within which most member species spend their entire evolutionary lifetimes (Hubbell 2003). Thus, we pooled estimates of species abundance within four ecoregions that belong to different biogeographic divisions to obtain the metacommunity SADs (Fig. 1, detailed in Appendix B). Estimates of biodiversity patterns in the ecoregions are summarized in Table 1.

**Fig. 1.**
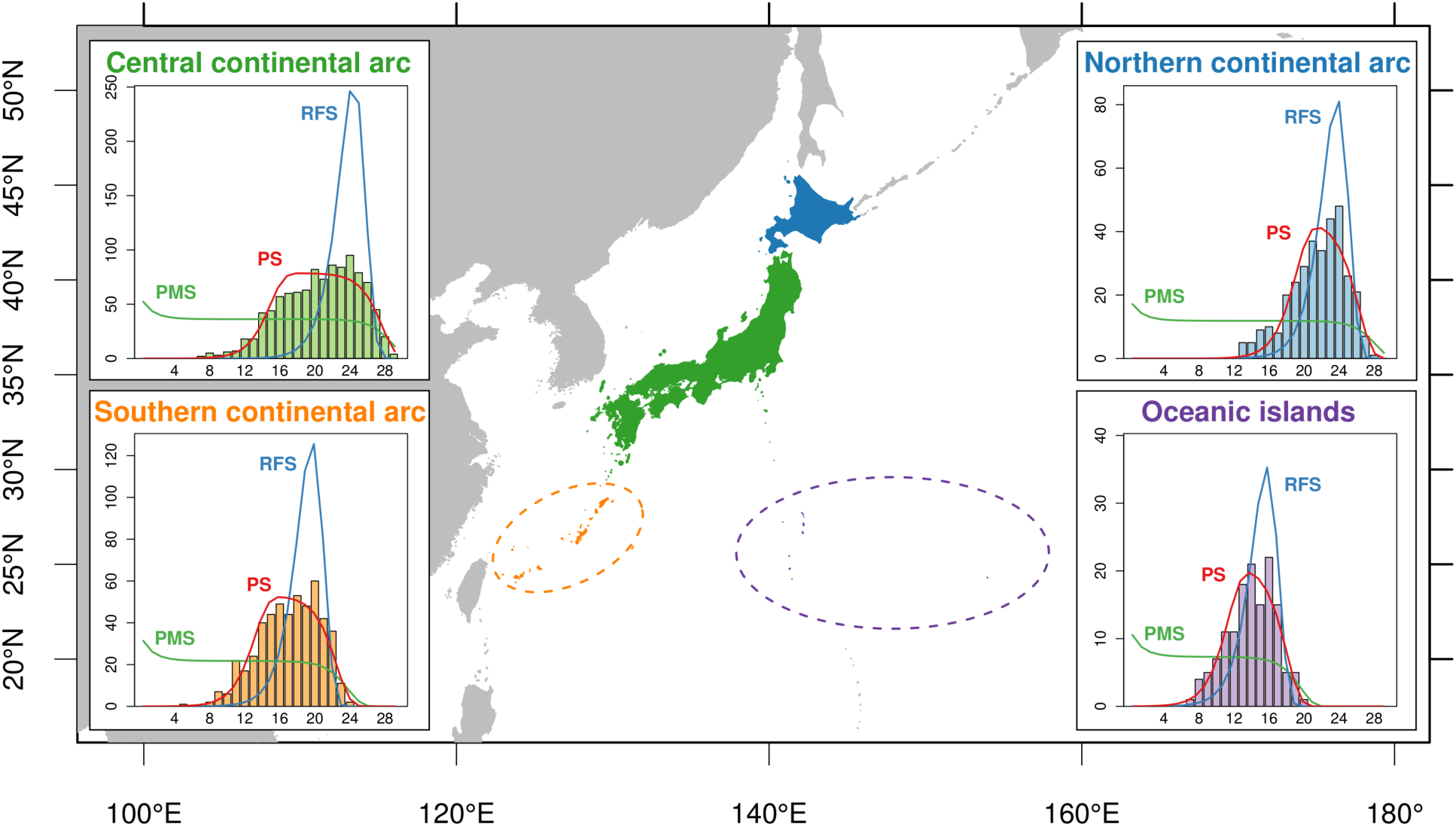
Metacommunity species abundance distribution in the four ecoregions of the East Asian islands. Ecoregions are discerned by colour (central continental arc: green, northern continental arc: blue, southern continental arc: orange, oceanic islands: purple). Histograms in the inner panels represent the estimated metacommunity species abundance distributions (SADs). The coloured lines represent metacommunity SADs predicted by the three variants of the unified neutral theory of biodiversity and biogeography (UNTB) (PMS – point mutation speciation model; RFS – random fission speciation model; PS – protracted speciation model) fitted to the metacommunity SADs. *x*- and *y*-axis indicate the abundance octave and number of species, respectively. The *j*th abundance octave is defined as the range of abundance *n* satisfying 2^*j*−1^ ≤ *n* < 2^*j*^.

**Table 1.** Estimates of community abundance, species richness, diversity index, and parameters relevant to the unified neutral theory of biodiversity. Species diversity is represented by Shannon entropy. Parameters related to the neutral models are: fundamental biodiversity number *θ*, speciation rate *ν*, and average species lifetime (generations) *L*.

|  | Ecoregion |  |  |  |
| --- | --- | --- | --- | --- |
|  | Central | Northern | Southern | Oceanic |
| Abundance |  |  |  |  |
| Total (metacommunity size $J_M$) | $1.67 \times 10^{10}$ | $3.37 \times 10^9$ | $3.29 \times 10^8$ | $6.35 \times 10^6$ |
| Mean | $4.73 \times 10^6$ | $3.40 \times 10^6$ | $2.27 \times 10^6$ | $3.53 \times 10^5$ |
| SD | $5.74 \times 10^6$ | $6.46 \times 10^6$ | $3.60 \times 10^6$ | $6.54 \times 10^5$ |
| Species richness |  |  |  |  |
| Total (γ-diversity) | 1024 | 328 | 508 | 141 |
| Mean (α-diversity) | 241.7 | 96.9 | 198.4 | 47.8 |
| SD | 75.7 | 33.0 | 81.1 | 34.2 |
| Shannon entropy |  |  |  |  |
| Total (γ-diversity) | 5.55 | 4.83 | 5.11 | 3.97 |
| Mean (α-diversity) | 4.58 | 4.13 | 4.18 | 2.66 |
| SD | 0.66 | 0.45 | 1.03 | 1.12 |
| Point mutation speciation model |  |  |  |  |
| $\theta$ | 52.27 | 17.15 | 31.40 | 10.56 |
| $\nu$ | $3.13 \times 10^{-9}$ | $5.09 \times 10^{-9}$ | $9.55 \times 10^{-8}$ | $1.66 \times 10^{-6}$ |
| $L$ | 19.6 | 19.1 | 16.2 | 13.3 |
| Random fission speciation model |  |  |  |  |
| $\theta$ | 1023.75 | 327.75 | 507.75 | 140.75 |
| $\nu$ | $3.76 \times 10^{-15}$ | $9.46 \times 10^{-15}$ | $2.38 \times 10^{-12}$ | $4.91 \times 10^{-10}$ |
| $L$ | $1.63 \times 10^7$ | $1.03 \times 10^7$ | $6.48 \times 10^5$ | $4.51 \times 10^4$ |
| Protracted speciation model |  |  |  |  |
| $\theta$ | 113.51 | 62.87 | 76.65 | 30.40 |
| $\nu$ | $2.65 \times 10^{-13}$ | $4.19 \times 10^{-14}$ | $2.96 \times 10^{-11}$ | $2.00 \times 10^{-9}$ |
| $L$ | $2.22 \times 10^5$ | $2.13 \times 10^6$ | $4.96 \times 10^4$ | $1.07 \times 10^4$ |

The SADs of metacommunities in the four ecoregions followed a left-skewed, lognormal-like distribution, whose short left tail indicates that the number of very rare species was negligible (Fig. 1). This pattern of the metacommunity SADs were consistently well explained by the protracted speciation model (Table 2). Point mutation speciation model fitted relatively well at the largest abundance classes, but failed to predict the number of less common species and rare species.

**Table 2.**
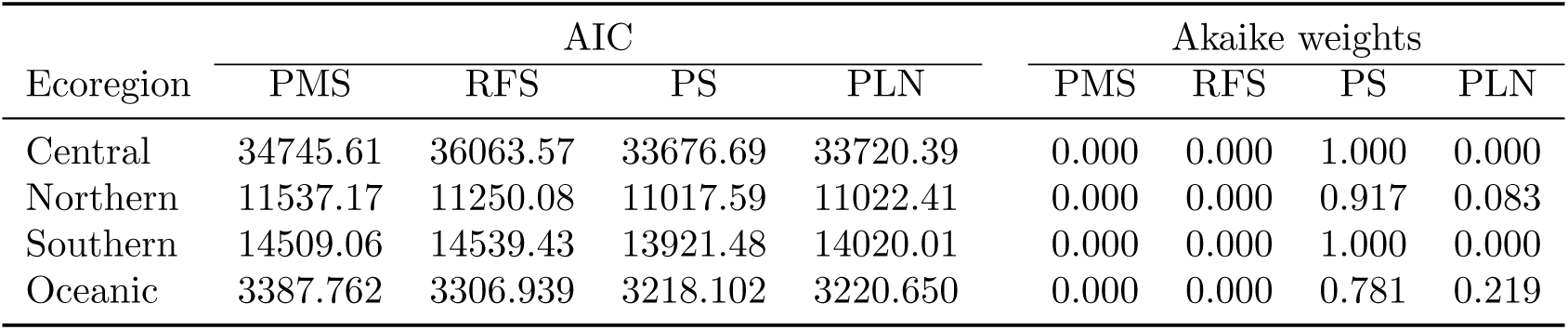
Model comparison for the fit of three variants of the unified neutral theory of biodiversity and biogeography (UNTB) and a Poisson lognormal model. Models were compared based on their “composite likelihood” suggested by Alonso & McKane (2004): see Appendix B for details on the procedures for model fitting and comparison. Abbreviations: PMS - point mutation speciation model; RFS – random fission speciation model; PS – protracted speciation model; PLN – Poisson lognormal model; AIC – Akaike information criterion.

Random fission speciation model overpredicted the number of moderately abundant species, while underpredicting the number of less common species. The results suggest that the manner of species diversification in these metacommunities was represented by neither of the two extreme modes, point mutation speciation or random fission speciation, but by an intermediate process expressed as a protracted speciation.

The macroscale SADs yielded estimates of the metacommunity size *J_M_* for each ecoregion, which enabled us to disentangle speciation rate *ν* from the fundamental biodiversity number *θ* (Table 1). A higher speciation rate and shorter average lifetime of a species was observed in ecoregions composed of small and isolated islands, the oceanic islands region, and the southern continental arc region (Table 1), implying relatively rapid evolutionary turnover of the metacommunity in those regions. The magnitude of *L* largely differed between the models; the point mutation speciation model predicted an average species lifetime of less than 20 generations, while the random fission speciation model predicted a very long lifetime, up to tens of millions of generations. Assuming that the average generation time of woody plants is about 30 years (Leigh *et al*. 1993, Nee 2005), the estimates of lifetime (i.e. hundreds of years in the point mutation speciation model and up to hundreds of millions of years in the random fission speciation model) are ecologically unrealistic for species. In contrast, the protracted speciation model provided moderate estimates of *L* that range from hundreds of thousands of years to tens of millions of years, which are comparatively congruent with previous estimates for species lifetime of vascular land plants based on fossil records (Niklas *et al*. 1983, 1985).

The UNTB, originally formulated with the point mutation and random fission speciation (Hubbell 2001), can fit well to empirical SADs at local communities. However, it has been criticized because of failing to explain the evolutionary aspects such as average species lifetime (Ricklefs 2003, Nee 2005, Ricklefs 2006). The concept of the protracted speciation achieved a considerable advancement of the UNTB and led to realistic predictions about macroevolutionary patterns of communities (Rosindell *et al*. 2010, Rosindell & Phillimore 2011, Etienne & Rosindell 2012). Nevertheless, in the explanation of empirical SADs, its superiority over the other speciation modes has been unapparent, probably due to limited sample size (Rosindell et al. 2010). Our study fulfils the gap between these theoretical and empirical developments in the UNTB by revealing metacommunity SADs across the four ecoregions in East Asian islands and provides a strong support for the protracted speciation model.

An analysis of metacommunity SADs also highlighted region-specific evolutionary processes, which can shape large-scale biodiversity patterns relevant to geographic characteristics (e.g. area, degree of isolation, and other physiographical conditions) of the regions (Qian & Ricklefs 2000, Xiang *et al*. 2004, Qian *et al*. 2017). Greater estimates of the speciation rate in regions of southern continental arc and oceanic islands than in the other two continental arc regions (Table 1) clearly indicate that these regions bear greater species diversity relative to their small land area (i.e. the metacommunity size). They are likely to reflect adaptive/non-adaptive radiation driven by historical vicariance (Kubota *et al*. 2014, 2017), which may have led these regions to act as “cradles of biodiversity” (Rangel *et al*. 2018). A fundamental limitation in our analysis was, however, that an immigration of new species realized by a long-distance dispersal from other biogeographic regions cannot be distinguished from an endemic diversification of species, and therefore the estimates of speciation rate represent the joint consequence of these two processes. Long-distance dispersal is another critical macroecological process (Jabot *et al*. 2008, Rosindell *et al*. 2011, Whittaker *et al*. 2017) which is especially likely to be promoted in the southern continental arc region by the repeated land bridge connections throughout the Cenozoic. Future studies exploring a further theoretical and methodological development to infer the relative role of speciation and long-distance dispersal are warranted (Etienne & Haegeman 2011).

The key element of the present study was the methodological development of an estimation of macroscale SADs that have been the inaccessible property of biodiversity in evolutionary ecology. Macroscale SADs indicate fundamental properties of the species pool such as the absolute size of communities and species abundance. Their accurate estimates are critically informative for both basic and applied field of ecology and biogeography; the proposed approach will improve the identification of the species pool (*γ* diversity) along geographical gradients (de Bello *et al*. 2012, Karger *et al*. 2016), facilitating our understanding of the origin and maintenance of biodiversity from an evolutionary perspective, the evaluation of the role of macroevolutionary processes (e.g. abiotic filtering and adaptive radiation) in community assembly, and the design of the protected areas network to capture biodiversity processes.

## Acknowledgements

We thank S. Eguchi and O. Komori for their helpful comments and discussion. We are grateful to T. J. Matthews for valuable comments and editing. We are particularly grateful to local botanists, vegetation researchers, and naturalists who have accumulated the information on plant distribution through their fieldwork steadies over the past decades. This research was supported by an allocation of computing resources of the SGI ICE X and SGI UV 2000 supercomputers from the Institute of Statistical Mathematics. Financial support was provided by the Japan Society for the Promotion of Science (no. 15H04424), the Environment Research and Technology Development fund of the Ministry of the Environment, Japan (4-1501), and Program for Advancing Strategic International Networks to Accelerate the Circulation of Talented Researchers, Japan Society for the Promotion of Science.

## Author Contributions

Y.K. conceived the ideas; B.K. and T.S. compiled the data; K.F. designed the methodology and conducted data analyses; J.F. contributed to data interpretion and model development; K.F. and Y.K. coordinated the writing of the manuscript. All authors discussed the results and contributed critically to the drafts.

## Methods

We developed a novel class of hierarchical models that can estimate SADs in discrete geographical units (i.e. grid cells) from spatially replicated multispecies detection-nondetection observations, in combination with various sources of data about the geographic distribution of species. The proposed model includes indicators of species presence and conditional individual density as its latent state variable, thereby enabling us to make an explicit prediction about the abundance of each species in each grid by fitting the model to available data. The formulation and statistical inference of the model are detailed in Appendix A.

The model was applied to a dataset of woody plant communities in midlatitude forests in Japan. The details of this application are fully described in Appendix B. Briefly, a large dataset comprised of 40,547 vegetation survey records collected within natural forests, species occurrence records, species distribution maps, and regional species checklists were used to estimate the abundance of 1,248 woody plant species within 4,684 ten-kilometre grid cells, which covered almost all the woody plant species and the entire land area of Japan. The estimates of species abundance, obtained through the empirical Bayes procedure, were then validated based on independent local abundance datasets of woody plant communities obtained in forest inventory plots. Although there was a tendency of underprediction, this validation has confirmed a positive correlation between the predicted and observed log abundance of woody plant species (Appendix B). It was also shown that the magnitude of the estimates of total abundance of woody plants in natural forests in the region was consistent with a recent global estimate of tree abundance (Crowther *et al*. 2015) (Appendix B).

Based on the results of model fitting, metacommunity SADs were obtained for the four ecoregions on the East Asian islands (i.e. the central, northern, southern, and oceanic region) by aggregating abundance estimates over grids within each region (Appendix B). For each ecoregion, three variants of the UNTB were fitted to the estimate of the metacommunity SAD. The fitted model included the point mutation speciation model (Hubbell 2001, Etienne & Alonso 2005), random fission speciation model (Etienne & Haegeman 2011), and protracted speciation model (Rosindell *et al*. 2010); for these models, a probability function of the metacommunity species abundance vector (i.e. likelihood function for metacommunity SAD) and/or an analytical solution of the SAD in the stationary metacommunity has been obtained and can be used for model fitting. Estimates of the speciation rate *ν* and mean species lifetime *L* were derived as a function of the estimated parameters (including *θ*) and metacommunity size *J_M_*.

### Data availability

The datasets generated and analysed during the current study are available from the corresponding author upon reasonable request.

## Appendix A: Statistical framework to estimate macroscale SADs

In this section, we describe a class of hierarchical models that estimates SADs in discrete geographical units (i.e. grid cells) from spatially replicated multispecies detection-nondetection observations, in combination with various data sources indicating the geographic distribution of species (Fig. 2). A hierarchical model is composed of a series of submodels, including an observation model describing the distribution of data conditional on some latent state variables and a system model describing the variation in the state variables (Royle & Dorazio 2008, Kery & Schaub 2012, Kéry & Royle 2016). In the following, we first describe a generalized linear mixed model (GLMM), which explains the multispecies detection-nondetection observations in terms of individual density of each species and therefore explicitly links binary observations to underlying SADs. Then, we extend this model to incorporate other sources of information about species occurrence that facilitate the inference of abundance for a number of species over a large geographical extent.

**Fig. 2.**
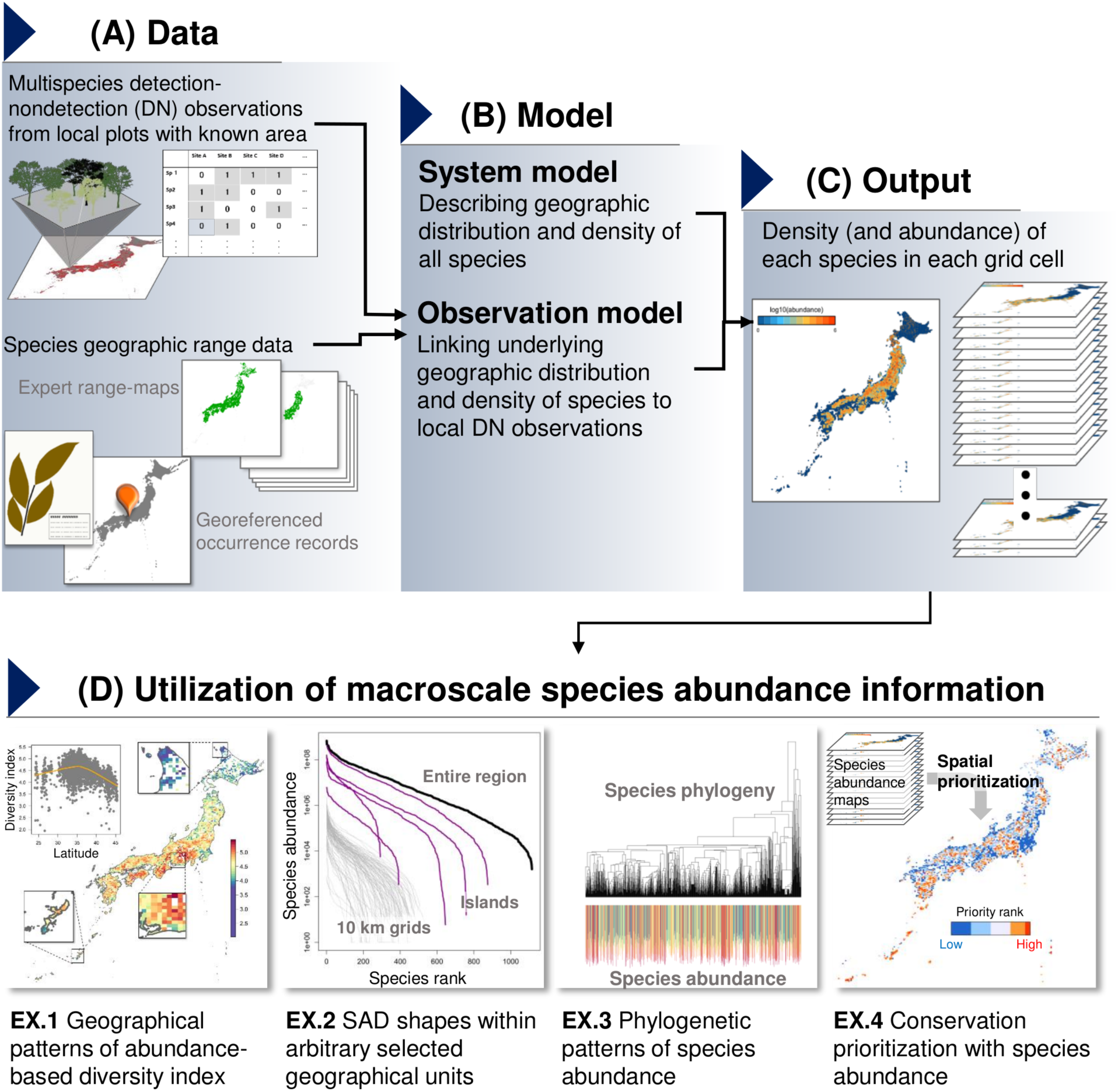
A framework for estimation of macroscale species abundance distributions (SADs). Spatially replicated detection-nondetection observations and various information on species geographic distribution (A) are integrated in a hierarchical model that links binary observations to underlying species abundance (B). A model fitting yields estimates of individual density of each species in each geographic grid, which can then be used to derive estimates of species abundance with the area of suitable habitat (C). The results can be used for diverse purposes relevant to e.g. community ecology, macroecology, biogeography, and applied fields of ecology (D).

### A model for spatially replicated detection-nondetection observations

We assume that there is a set of geographic areas of interest that contain *I* species of interest and are divided into *J* geographical grids. Suppose that grid *j* (*j* = 1,…, *J*) contains *K_j_* > 0 replicated sampling plots in which occurrence was assessed for each species. We denote detection (1) or nondetection (0) of species *i* in plot *k* in grid *j* as *y_ijk_* (*i* = 1, …, *I*; *j* = 1, …, *J*; *k* = 1,…, *K_j_*). We also assume that the area of each sampling plot was recorded, and denote the area of sampling plot *k* in grid *j* as *a_jk_*.

The goal of the inference is to estimate the abundance of each species within each grid from these locally replicated detection-nondetection observations. To achieve this, we explicitly make several key assumptions in the data generating process. First, we assume that individuals are distributed within some suitable habitats (e.g. forests) in which sampling plots are placed so that they never overlap. Second, we assume that for each grid the spatial point pattern of individuals within the habitats can be regarded as an independent superposition of homogeneous Poisson point processes, each of which represents the spatial alignment of individuals of a species. In the ecological context, this assumption implies that the centres of individuals are regarded as points, and individuals are distributed independently of one another with species-specific individual densities that are constant within a grid (Illian *et al*. 2008).

These assumptions give us a probability function that explicitly links the probability of species detection within a plot to the density of that species in the grid. Let us denote the individual density of species *i* in grid *j* by *d_ij_*. Then, the number of individuals occurring in a plot of area *a_jk_* independently follows a Poisson distribution with a mean of *d_ij_a_jk_* (Illian *et al*. 2008). Therefore, the probability for detecting at least one individual of species *i* in plot *k* in grid *j*, *p_ij_a_jk_*, can be written as:

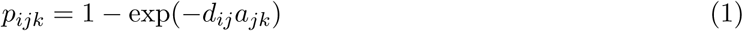

where exp(–*d_ij_a_jk_*) corresponds to the probability mass of a Poisson distribution with a mean *d_ij_a_jk_* at zero (i.e. a probability that the plot captures no individuals).

On the basis of these settings and assumptions, we provide a state space formulation of the first hierarchical model we consider, in which the model is described in terms of a series of submodels that are conditional on latent state variables and parameters (Royle & Dorazio 2008, Kéry & Schaub 2012, Kéry & Royle 2016). The latent variable of the model was the grid-level individual density of species, which we have already defined as *d_ij_*.

The observation model describes the occurrence of species within a sampling plot. We can regard the detection-nondetection observation of species, *y_ijk_*, as a random variable that independently follows a Bernoulli distribution with a detection probability *p_ijk_*:

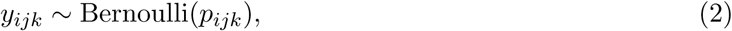

where *p_ijk_* is determined by Equation (1) under the assumption of the superposed homogenous Poisson point process.

The system model describes variation in the individual density *d_ij_*. We decompose the logarithm of *d_ij_* into an intercept term *μ* and three normally distributed random effects, species
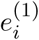,
grid
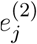, and the combination of species and grid
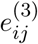:

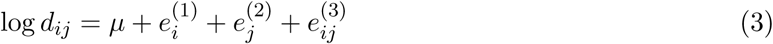

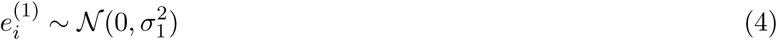

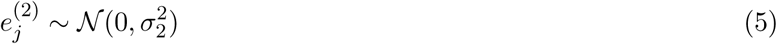

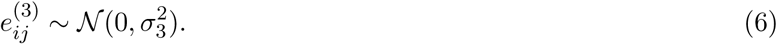

These submodels jointly construct a Bernoulli GLMM with complementary log-log link, in which *a_jk_* is treated as an offset term. The model can therefore be fitted to data with standard GLMM packages that implement multiple random effects, such as **lme4** in **R** (Bates *et al*. 2015).

The model described above has a relatively simple structure, in which variation in individual density was explained only by several unstructured random effect components. The inclusion of random effects is essential in a multispecies distribution modelling as it enables us to “borrow strength” in the inference: it will improve the estimates for grids with few replicated plots and/or the estimates for rare species because information is shared across all grids and species through common distributions specified for random effects (Iknayan *et al*. 2014, Warton *et al*. 2015, Evans *et al*. 2016). In an analogous fashion to many other classes of hierarchical models and species distribution models (SDMs), environmental covariates could also be introduced in the system model to explicitly describe the association between environmental factors and individual density. In addition, the model could also explain the correlation structure of random effects on the geographic and/or phylogenetic space in an explicit manner (Ives & Helmus 2011, Kaldhusdal *et al*. 2015). Such generalizations will potentially enhance the model prediction and provide further ecological insights. However, they may be difficult to adopt in practice, especially in studies that examine a very large number of species and grids, as is the case with our application described in Appendix B, because the model may involve an excessive number of parameters and/or a huge covariance matrix, rendering the inference computationally challenging (Warton *et al*. 2015).

### Integrating grid-level occurrence information

Owing to the fact that information is shared by random effects, the simple random effect model without any covariate can still provide estimates of individual density that are specific to each species and grid. However, the estimates may be inaccurate especially in grids where the number of plots is limited and species density is low. To overcome this issue, we extend the model to integrate replicated detection-nondetection observations with data that may directly inform about the grid-level presence-absence of species such as species occurrence records and expert range maps.

We introduce a latent indicator state variable that represents the grid-level presence-absence of species and is denoted as *z_ij_*. The detection probability *p_ijk_* is then expressed as follows:

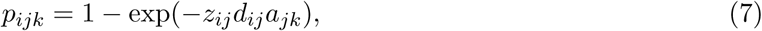

which indicates that the detection probability is 0 when the species is absent in the grid (*z_ij_*=0) but it takes 1 – exp(–*d_ij_a_jk_*) when the species is present in the grid (*z_ij_* = 1). Hence, *d_ij_* now represents the individual density that is *conditional* on the presence of that species.

We regard *z_ij_* as a random variable following a Bernoulli distribution and add an additional system model component to describe it. By adopting a similar modelling approach applied for the individual density, the additional components can be constructed as follows:

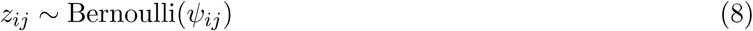

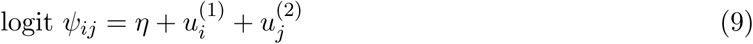

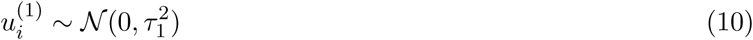

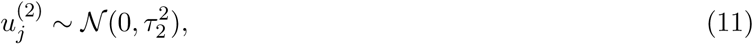

where *ψ_ij_* is the occurrence probability of species *i* in grid *j*, which was decomposed into an intercept term *η* and two normally distributed random effects that vary over species
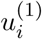
and grids
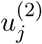
on a logit scale.

We assume that the grid-level species occurrence *z_ij_* is partially observed via the plot-level detection-nondetection observations and/or the auxiliary grid-level presence-absence information. A grid-level presence of species may be registered, for example, by museum- or herbarium-based specimens and/or occurrence records, while absence of species may be deduced by exploiting, for example, expert range maps (Merow *et al*. 2017) and/or regional species checklists. In general, the information about the species absence should be treated conservatively because it is difficult to verify (Merow *et al*. 2017); therefore, a larger weight should be placed on the evidence of species presence than on that of species absence if different sources of data are in conflict.

Under these considerations, the conditional likelihood defined by our observation model (Equation 2) takes two cases depending on whether the presence-absence of the species is known or not. Formally, we denote the vector of all parameters (i.e. *η*, *μ*, *τ*_1_,*τ*_2_,*σ*_1_,*σ*_2_,*σ*_3_) and the vector of all random effects
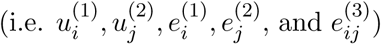
by ***θ*** and ***ξ***, respectively. Let *x_ij_* = 1 denotes that *z_ij_* is known for species *i* in grid *j* and *x_ij_* = 0 denotes otherwise. Then, by letting **y**_*ij*_ = (*y*_*ij*1_,…, *y_ijK_j__*) and **D**_*ij*_ = (**y**_*ij*_,*z_ij_*), the conditional likelihood, *p*(**D**_*ij*_ | ***ξ***, ***θ***), can be expressed as follows:

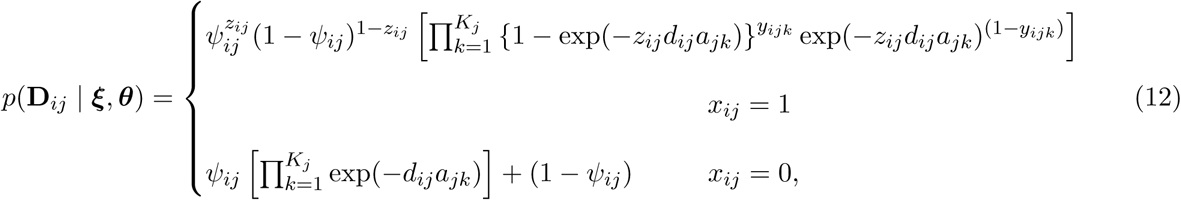

where in the former case, the conditional likelihood is given as a joint likelihood of **y**_*ij*_ and *z_ij_*, and in the latter case, it is given by the marginalized likelihood of **y**_*ij*_ because *z_ij_* is missing. We note that *d_ij_* and *ψ_ij_* are respectively a function of ***ξ*** and ***θ*** (Equations (3) and (9)), although that is not expressed explicitly in the right-hand side of the equations.

In this integrated model, geographical grids that contain no detection-nondetection observations but have grid-level presence-absence information for some species can still contribute to the inference of parameters. Let us now assume that the set of geographical areas of interest is divided into *J* geographical grids, in which grid *j* (*j* = 1,…,*J*) contains *K_j_* ≥ 0 plots. Then, for grid *j* such that *K_j_* > 0, the conditional likelihood is expressed by Equation (12), and for other grids (*K_j_* = 0), it is written as:

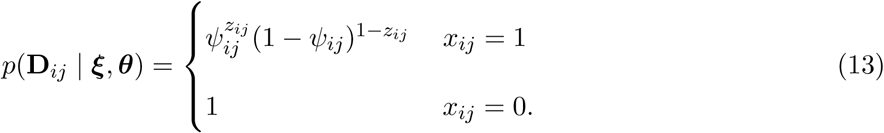

### Statistical inference

As a class of general hierarchical models, the integrated model can be fitted to data by using either maximum marginal likelihood (also known as empirical Bayes) or fully Bayesian approach. Let us denote **D** as the vector of all data. In both approaches, inference is based on a joint distribution of data and random effects, *p*(**D**, ***ξ*** |***θ***), which is also known as a complete data likelihood (King 2014). In the former approach, estimation can be achieved via a two-stage procedure, where parameters are estimated by maximizing a marginal likelihood *p*(**D**| ***θ***) = *∫ p*(**D**, ***ξ*** | ***θ***)*d****ξ*** and then, maximum a posteriori probability (MAP) estimates of random effects can be obtained conditionally on the parameter estimates
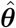
by maximizing *p*(**D**, ***ξ*** | 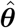). Although an evaluation of the marginal likelihood may be computationally challenging, some recently developed software, such as **AD Model Builder** (Fournier *et al*. 2012) and **Template Model Builder** (Kristensen *et al*. 2016), can efficiently approximate the marginal likelihood of a wide class of hierarchical models by using the Laplace approximation. In contrast, in the latter approach, the focus of inference is the joint posterior distribution of parameters and random effects *p*(***θ***, ***ξ*** |**D**) =
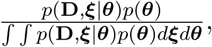
where a prior distribution for parameters *p*(***θ***) is needed to be specified. Although the integration over parameters and random effects is not tractable in general, a Markov chain Monte Carlo (MCMC) method can be used to obtain random samples from the posterior distribution. Several generic software are available to run MCMC for a vast array of hierarchical models (e.g. Plummer 2003, Carpenter *et al*. 2017).

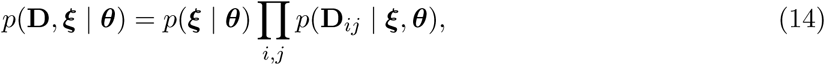

where *p*(**D**_*ij*_ | ***ξ***, ***θ***) is the conditional likelihood derived from the observation model (Equations 12–13), and *p*(***ξ*** | ***θ***) represents a probability density of random effects that is determined by the system model (Equations 4–6 and 10–11):

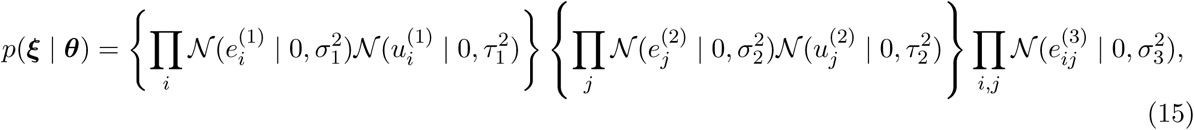

where 𝓝(*x* | 0, *σ*^2^) denotes the probability density of a normal distribution with mean 0 and variance *σ*^2^ evaluated at *x*.

Once estimates (or posterior samples, in case of fully Bayesian approach) of random effects are obtained, we can derive the estimates of *ψ_ij_* and *d_ij_*, denoted by
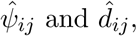
respectively, by substituting the estimates of random effects into Equations (3) and (9), respectively. Based on these estimates, we can further derive estimates for a wide array of variables that are of ecological interest. For example, the number of modelled species, denoted by *S_j_*, that are actually present in grid *j* can be estimated as:

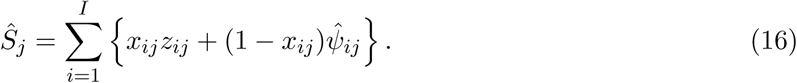

Note that the use of the estimated occurrence probabilities
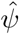
enables this estimator to account for the possibility of the presence of species even when they are not detected in the replicated plots (*c.f*., Dorazio & Royle 2005) or no detection-nondetection observation is available in the grid. Let **N_*j*_** denotes the vector of abundance of all species in grid *j*. This vector represents the SAD, the property of an ecological community that we aimed to infer, and can be estimated for each grid as:

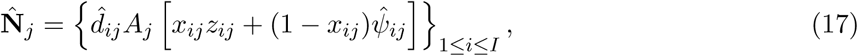

where *A_j_* denotes the area of habitats in grid *j*. We can also estimate the SAD for a subset of the area of interest *𝓘*, denoted by 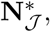 as follows:

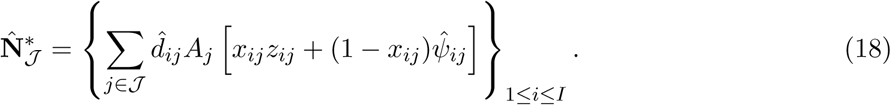

Note that the estimates of abundance of each species further permit to obtain various diversity indices that are a function of a vector of (relative) abundance, such as Shannon entropy and Gini-Simpson index, as well as other generalized metrics including phylogenetic/functional diversity indices and the Hill numbers (Chao *et al*. 2014).

### Related models

Related classes of models that motivated our method include the Royle-Nichols model, which estimates the abundance of animals that are not detected perfectly from spatially replicated detection-nondetection observations (Royle & Nichols 2003), and its extension to community data developed by Yamaura *et al*. (2011). However, the proposed model may appear largely different from these models because both observation and system process are modelled differently: the models are rather aimed to describe observations of mobile animals that are subject to imperfect detection and thus do not assume Poisson point processes to derive an observation model. Another closely related class of models is the multispecies site occupancy model which explains detection-nondetection observations of a number of species simultaneously in terms of the occurrence of species at each site (Dorazio & Royle 2005, Dorazio *et al*. 2006). Indeed, our estimator for species richness (Equation 16) resembles that derived in Dorazio & Royle (2005). Fithian *et al*. (2015) introduced a multispecies version of the species distribution model (SDM) which integrates presence-absence data into the inhomogeneous Poisson process model for presence-only data. Their model component for presence-absence observations is a Bernoulli generalized linear model (GLM) with complementary log-log link (see also the related discussion by Dorazio (2014)). Models that jointly infer geographical distribution of many species have been recently named the joint species distribution models (JSDMs) (Warton *et al*. 2015).

## Appendix B: An application to woody plant communities in East Asian islands

### Estimation of species abundance and its validation

We applied the proposed model to a dataset of woody plant communities in midlatitude forests in Japan. For the replicated detection-nondetection observations, we compiled a large dataset from vegetation surveys that consists of 40,547 georeferenced plots placed in natural forests between 24°02′–45°30′ N and 122°56′–153°59′ E, which comprises the dataset of Kusumoto *et al*. (2015) and the national vegetation survey of Japan (http://www.biodic.go.jp/english/kiso/vg/vg_kiso_e.html). The plot area ranged from 0.01 m^2^ to 18,000 m^2^.

In the vegetation survey, species occurrence in the sampling plots (called “relevés”) is traditionally recorded according to cover classes for individual species. We converted these vegetation observations into detection-nondetection records by assigning 1 if the species appeared in the plot and 0 otherwise. In this analysis, we standardized the names of woody plant species and pooled the data for varieties and subspecies with those of their parent species. As a result, we obtained detection-nondetection observations for 1,248 species, which covers almost every woody plant species found in Japan. We divided the entire study area into 10 × 10 km grids (Kubota *et al*. 2015, 2017). We analysed in total 4,684 grids which covered ca. 99.5 % of the total land area of Japan. In total 3,695 grids contained at least one vegetation plot.

We also compiled the species occurrence information at a grid level based on multiple data sources. Species presence was registered from museum and herbarium specimens, species occurrence records, and distribution maps of plant species compiled in Horikawa (1972). Species absence was recorded from the distribution maps of Horikawa (1972) and regional species checklists compiled by prefectures of Japan.

The integrated model was fitted to these data by using the empirical Bayes estimation procedure implemented in the **Template Model Builder** (Kristensen *et al*. 2016), with the aid of **TMB** package (version 1.7.10) run in **R** (version 3.2.0). The estimates (and standard errors) of parameters were:
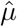 = 4.575 (0.043),
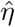 = −3.267 (0.074),
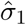 = 1.373 (0.030),
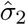
= 0.680 (0.009),
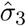 = 1.217 (0.002),
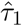 = 2.551 (0.052), and
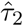 = 0.956 (0.010).

Based on the model estimates, the abundance of 1,248 woody plant species within natural forests was estimated for 4,684 grids by using Equation (17). The area of natural forest in each grid was obtained based on the national survey of the natural environment (http://www.biodic.go.jp/trialSystem/EN/info/vg.html).

The total woody plant abundance within the natural forest in Japan was estimated to approximately 20.4 billion, with the abundance of individual species ranging over six orders of magnitude, from species with 10^8^ individuals to species with hundreds of individuals. The estimated total abundance approximately corresponded to 0.671% of the recent estimate for the number of trees worldwide (3.04 trillion; Crowther *et al*. 2015). This percentage parallels that of the total area under natural forests in Japan (0.367%) in relation to the area of forests around the globe, which was calculated based on the FAO statistics for 2015. Therefore, our estimate seems largely consistent with the global estimate of tree abundance (Crowther *et al*. 2015), which was independently obtained by using entirely different datasets and inference approaches.

The result highlighted geographical and latitudinal patterns of biodiversity over the East Asian islands (Fig. 3). The total abundance of woody plants revealed no apparent distinct latitudinal patterns, although it tended to be slightly smaller at lower latitudes where few large islands exist (Fig. 3A). By contrast, species richness and diversity index (represented by Shannon entropy) exhibited a clear, and similar, hump-shaped latitudinal gradient: species diversity was highest in the midlatitude zone of the Japanese archipelago, which has a substantial amount of land area, and decreased in both north and south directions (Fig. 3B, C). We observed that compared to species richness, diversity index shows a more mosaic-like geographical pattern (Fig. 3C). Estimates of species richness correlated strongly (Pearson’s correlation coefficient 0.93; results not shown) with another set of estimates of species richness within 10 km square grid in the same region, which was obtained based on a different (while partially in common) dataset and inference (Kubota *et al*. 2015).

**Fig. 3.**
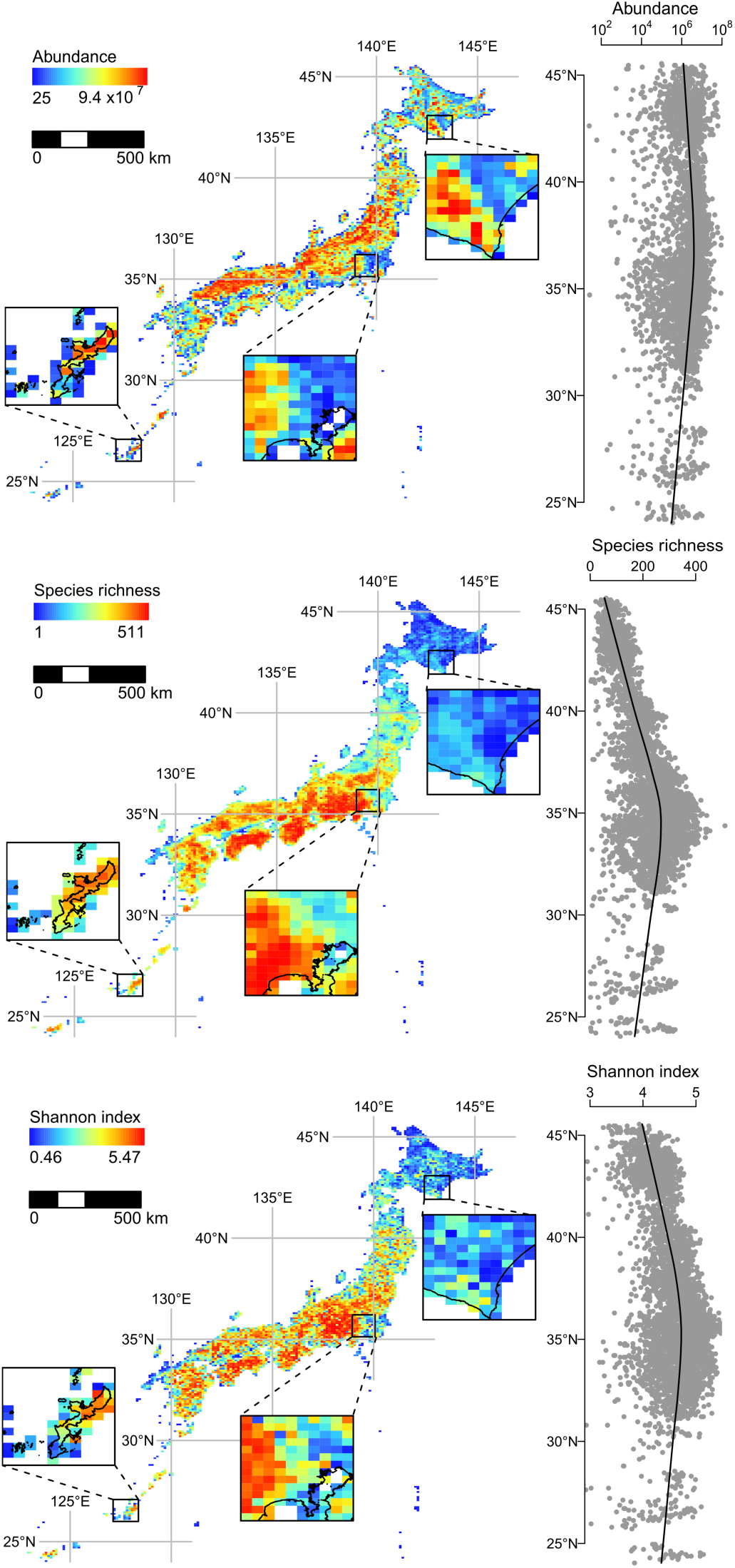
Maps of community properties estimated in 10 km square grids. (A) total number of individuals (abundance), (B) number of species (species richness) and (C) species diversity index (Shannon entropy). To illustrate finer spatial patterns, three arbitrarily selected sections are enlarged.

The estimates of species-specific abundance were validated based on data from geographically replicated forest inventory plots that were independent of the fitted data. We used three sources of forest inventory data that were collected in natural forests in Japan. They include the forest dynamics plots (FDP), the national forest inventory plots (NFI), and forest sampling plots along latitudinal and elevational gradients (FSLE). Sampling procedures and spatial coverage differed between the inventory data as we explain below.

The FDP dataset consists of species abundance data collected from 40 quadrats. In each quadrat, which was usually 1 ha in size, individuals with a diameter of ≥ 15 cm at breast height (DBH) were monitored (http://www.biodic.go.jp/moni1000/forest.html). This dataset fairly represents the mosaic structure of forests with different developmental stages and thus is expected to precisely capture local population size for common climax species in old growth mountain forests, while it may poorly represent the population of pioneer or fugitive species, especially in lowland forests.

The NFI dataset included 7,674 plots in which woody plant individuals were assessed in nested concentric circular plots. Individuals with DBH > 1 cm were measured in a 0.01 ha circular area, while those with DBH > 5 cm and > 18 cm were surveyed in a 0.04 ha and 0.1 ha circle, respectively (http://www.rinya.maff.go.jp/j/keikaku/tayouseichousa/). The NFI plots were systematically placed in a 4 km × 4 km grid laid over entire Japan and thus were expected to provide less-biased samples of density of woody plants.

The FSLE dataset included 460 plots where woody plant individuals were surveyed in a 0.01 ha area (unpublished data by Y. Kubota). Plots were placed along the elevational and latitudinal gradients and thus were expected to reflect the environmental heterogeneity in the midlatitude forests.

For each grid that contains at least one forest inventory plot, observed abundance was compared to predicted abundance that was derived based on the model estimates. In order to predict the abundance in NFI plots, we set the area of each plot to 0.1 ha.

The predicted and observed log abundance of woody plant species were mildly correlated and generally distributed around the identity line, although a tendency of the model to underpredict the abundance was also evident (Fig. 4). A possible explanation for this tendency of underprediction is the assumption of a superposed homogeneous Poisson point process for the spatial alignment of individuals, which was adopted to estimate the density of woody plants from replicated detection-nondetection observations (Appendix A). This assumption was indeed ecologically implausible, and may lead to an underestimation of individual density when violated because a spatial clustering of individuals inflates the probability of nondetection of species within a sampling plot (He & Gaston 2000, Yin & He 2014). We would therefore regard the model as giving a “first approximation” of species abundance in a large spatial extent. Although the model highlighted the geographical structure of biodiversity, a future modeling effort for accommodating more ecological realities are warranted to obtain better estimates.

**Fig. 4.**
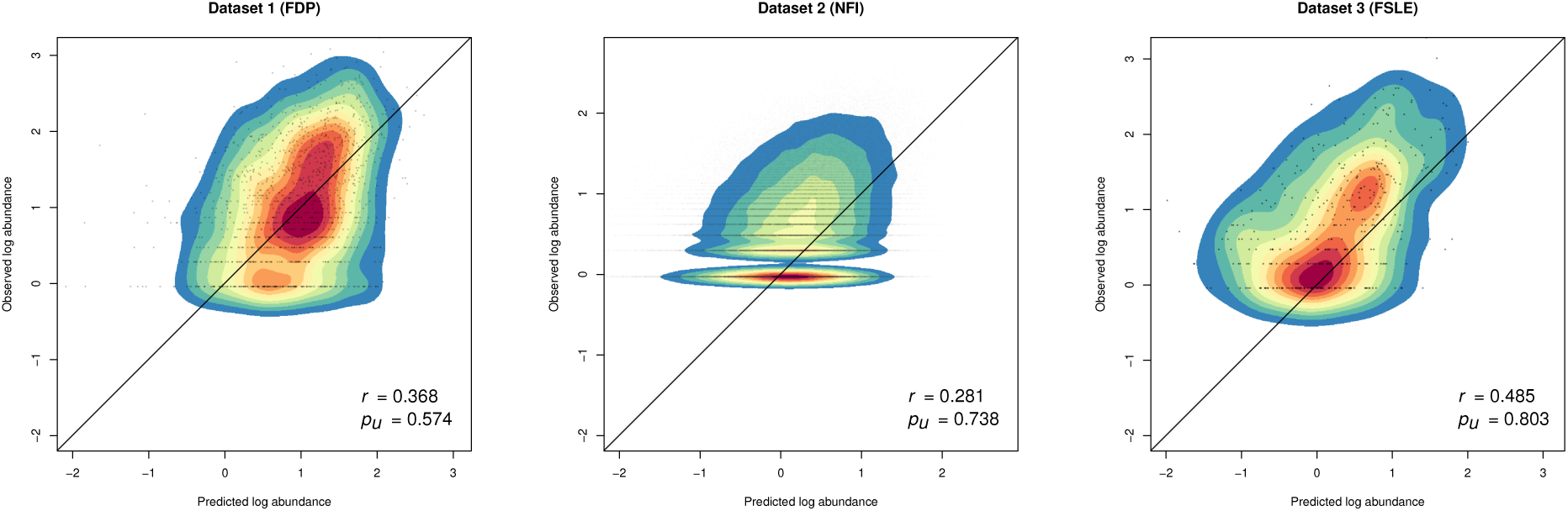
Result of the model validation. Pearson’s correlation coefficient (*r*) between observed log abundance and predicted log abundance of species, and the probability of underprediction (*p_u_* = Pr[Observed log abundance > Predicted log abundance]) are shown for three validation datasets of forest inventory plots, forest dynamics plots (FDP), national forest inventory (NFI), and forest sampling plots along latitudinal and elevational gradients (FSLE). The crossed lines are the identity lines.

### Inference of metacommunity SADs

Based on a previous biogeographic assessment of woody plants in the Japanese archipelago (Kubota *et al*. 2014) and Takhtajan’s floristic provinces (Takhtajan 1986), we divided the archipelago into four ecoregions (Fig. 1) and obtained metacommunity SADs by aggregating abundance estimates over grids within each ecoregion (Equation 18). The four ecoregions are defined as follows: (1) The *central continental arc region* is the largest ecoregion, which includes the three largest islands in Japan (Honshu, Shikoku, and Kyusyu). It encompasses deciduous and evergreen broad-leaved forests and belongs to the Takhtajan’s Japan-Korea province; 3,530 geographical grids belong to this ecoregion. (2) The *northern continental arc region* is the second largest ecoregion, and it includes Hokkaido, the second largest island of Japan. It encompasses coniferous and deciduous broad-leaved forests and belongs to the Takhtajan’s Sakhalin-Hokkaido province. The Tsugaru Strait separates the central continental arc region and northern continental arc region; 991 geographical grids belong to this ecoregion. (3) The *southern continental arc region* is composed of the Nansei Islands and separated from the central region by the Tokara Strait. It encompasses evergreen broad-leaved forests and belongs to the Takhtajan’s Tokara-Okinawa province. This ecoregion comprises 145 geographical grids. (4) The *oceanic islands region* is composed of the Bonin (Ogasawara) Islands. It encompasses evergreen broad-leaved forests and belongs to the Takhtajan’s Volcano-Bonin province. Differing from other ecoregions, in which almost all the lands are continental islands, the oceanic region is composed of oceanic islands only. It includes 18 geographical grids.

For each ecoregion, we fitted and compared three variants of the unified neutral theory of biodiversity and biogeography (UNTB) to the estimate of the metacommunity SAD. The fitted model includes the point mutation speciation model (Hubbell 2001, Etienne & Alonso 2005), the random fission speciation model (Etienne & Haegeman 2011), and the protracted speciation model (Rosindell *et al*. 2010); for these models, a probability function of the metacommunity species abundance vector (i.e. likelihood function for metacommunity SAD) and/or an analytical solution of the SAD in the stationary metacommunity has been obtained and can be used for model fitting.

The point mutation speciation model was fitted to the metacommunity SADs by using maximum likelihood method. The likelihood function for metacommunity SAD (i.e. assuming no dispersal limitation) under point mutation speciation model is known as the Ewens sampling formula (e.g. Equation 2 in Etienne & Alonso 2005). Formal likelihood-based inferences were, however, difficult to obtain for the other two models. Although a sampling formula has been acquired for a metacommunity under random fission models (Equation 38 in Etienne & Haegeman 2011), we were not able to apply this formula to our specific data as it underflows when the size of metacommunity is large, even when high precision arithmetic is used. We thus reached a compromise to use Equation 21 in Etienne & Haegeman (2011), which was derived without considering the sampling process, but provides the equilibrium probability function of the species abundance vector in a metacommunity with a fixed size *J_M_*. For protracted speciation model, no likelihood function was available. To fit this model, Rosindell *et al*. (2010) used “composite likelihood” that was suggested by Alonso & McKane (2004). This approach was however not practical in our case because of the large metacommunity size, thereby requiring an excess number of evaluations of the expected number of species with specific abundance. This prohibited its adoption in the numerical optimization procedure. We therefore applied a least square method to the Preston’s abundance octaves of metacommunities. We note that in addition to these three models, we also fitted the per-species speciation model of Etienne *et al*. (2007). However, this model consistently yielded boundary estimates that made the model identical to the point mutation speciation model. We thus omitted it from the comparison.

These differences in the fitting procedure render the model comparison complicated. To compare fitting of the models, while accounting for differences in the number of parameters (we note that point mutation model and random fission model have one free parameter (*θ*) but protracted speciation model has two (*θ* and *β*)), we prefer to use information criteria (McGill 2003, McGill *et al*. 2007) which relies on the formal maximum likelihood inference (Konishi & Kitagawa 2008). However, to fully utilize this approach was impossible in our application because the likelihood function was available only in the point mutation model. Thus, we compared the models based on the Akaike information criterion (AIC) and the Akaike weights (Burnham & Anderson 2002) calculated with “composite likelihood” (Alonso & McKane 2004), assuming that the parameter estimates of the random fission speciation model and protracted speciation model attain the maximum likelihood. In the model comparison, we also included a Poisson-lognormal mixture model (Bulmer 1974) as a flexible, simple baseline statistical model (McGill *et al*. 2007).

The objective function (i.e. negative log-likelihood or sum of squared error) of the variants of UNTB was minimized in terms of fundamental biodiversity number *θ* (in addition to *β*, in the case of protracted speciation model). Estimates of the speciation rate *ν* and mean species lifetime *L* were then derived as a function of these estimated parameters and metacommunity size *J_M_*. In point mutation model, *ν* relates to other quantities as
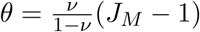
(Etienne & Alonso 2005), whereas in random fission model, the relationship is given as
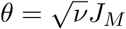
(Etienne & Haegeman 2011). In the protracted speciation model, the corresponding equation is given as
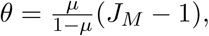 where
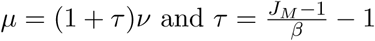
(Rosindell *et al*. 2010). The average species lifetime is obtained from the general equation of Ricklefs (2003):
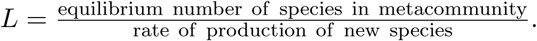
The corresponding formula is as follows: for point mutation speciation model, *L* ≈ – log *ν*; for random fission speciation model, *L* ≈
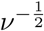
(Etienne & Haegeman 2011); for protracted speciation model, *L* ≈ – *τ* log *τμ* (Rosindell *et al*. 2010).

